# Time-resolved transcriptomic mapping reveals conserved stress programs and metabolic rewiring in *Escherichia coli* under antimicrobial photodynamic and blue light exposure

**DOI:** 10.64898/2026.02.02.703343

**Authors:** Natalia Burzyńska, Michał Wojciech Szcześniak, Mariusz Grinholc

## Abstract

Antimicrobial photodynamic inactivation (aPDI) and antimicrobial blue light (aBL) are emerging, resistance-agnostic strategies for controlling bacterial pathogens, yet their systems-level impact on prokaryotic physiology remains incompletely understood. Here, we used time-resolved global transcriptomics to define how *Escherichia coli* reprograms gene expression in response to diverse photodynamic stresses. *E. coli* K-12 BW25113 was exposed to five phototreatments differing in photosensitizer chemistry and light wavelength, including rose bengal, TMPyP, new methylene blue, aBL alone, and aBL combined with 5-aminolevulinic acid, and transcriptional responses were profiled after short (30 min) and prolonged (7–8 h) exposure. Short-term phototreatments triggered rapid and extensive transcriptional remodeling, affecting up to ∼58% of the genes and dominated by conserved stress programs including oxidative defense, sulfur metabolism, and a broad downshift in biosynthesis and energy generation. In contrast, prolonged exposure elicited more restrained but highly treatment-specific adaptive responses, characterized by suppression of core energy metabolism, including oxidative phosphorylation and the tricarboxylic acid cycle, coupled with activation of alternative catabolic pathways. Together, these findings reveal a common acute stress architecture across photodynamic modalities followed by divergent long-term adaptive trajectories, providing a systems-level framework for understanding bacterial responses to light-based antimicrobials and informing the rational optimization of photodynamic therapies.

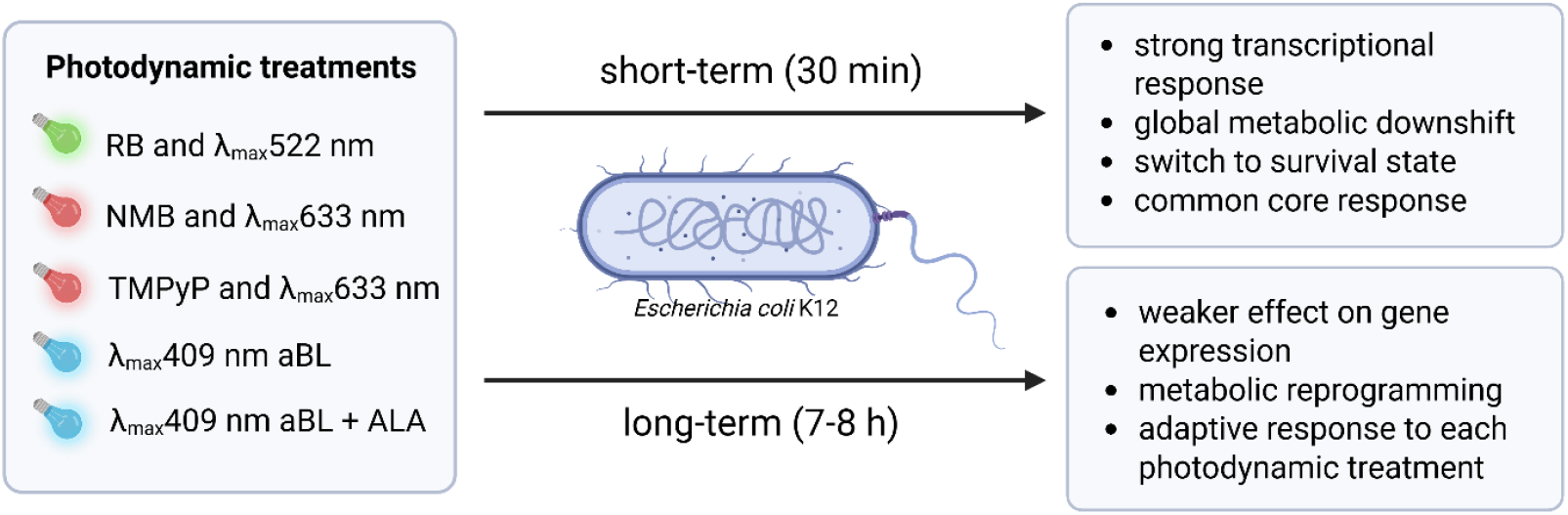

## Introduction

The growing crisis of antimicrobial resistance (AMR) has emerged as one of the most pressing global health challenges, threatening not only human and animal health but also food production systems and environmental sustainability. In response, the World Health Organization (WHO) has called for a coordinated, multisectoral strategy to mitigate AMR under the One Health initiative, which emphasizes the cooperation of human, animal, and environmental health sectors [1].

Antimicrobial photodynamic inactivation (aPDI) and antimicrobial blue light (aBL) align with this initiative, offering environmentally friendly, non-antibiotic approaches capable of eradicating pathogens regardless of their antimicrobial susceptibility profile, without inducing resistance [2], [3], [4]. The principal mechanism involves a light, non-toxic photosensitizer (PS) and molecular oxygen to induce an oxidative burst that non-selectively kills bacteria. While both methods share this general mode of action, they differ primarily in the origin of the photosensitizers involved. In aPDI, exogenous photosensitizing compounds are introduced and activated by specific light wavelengths matching their absorption spectra [5]. In contrast, aBL relies on the excitation of endogenous porphyrins naturally present within bacterial cells, which absorb blue light in the 400–470 nm range [6]. Following illumination, both exogenous and endogenous PSs reach a long-lived triplet state and can react by three proposed mechanisms. In the Type I mechanism, electron transfer leads to the formation of superoxide radicals (0_2_^-•^), hydrogen peroxide (H_2_0_2_), and hydroxyl radicals (HO•). The Type II mechanism results in energy transfer directly to molecular oxygen, yielding highly reactive singlet molecular oxygen (^1^O_2_) [7]. These ROS (reactive oxygen species) can cause oxidative damage to bacterial membranes, nucleic acids, and essential biomolecules, ultimately leading to bacterial cell death [8], [9]. Additionally, a Type III mechanism has been proposed, in which light-activated photosensitizers exert bactericidal effects through direct interaction with cellular targets, independently of oxygen availability [10].

Despite their proven antimicrobial efficacy, both aPDI and aBL remain underutilized in clinical practice, largely due to the lack of standardized protocols and insufficient understanding of bacterial response mechanisms. The photodynamic treatment strategies can vary widely in photosensitizers, light sources, and treatment durations; this heterogeneity complicates clinical translation. To date, only a few studies have addressed global transcriptomic or regulatory responses following aPDI or aBL treatment [11], [12], [13], [14]; these have typically been limited to a single photosensitizer or specific time point, without systematic comparison across multiple treatments or exposure durations.

To comprehensively elucidate the molecular responses of *Escherichia coli* to light-based antimicrobial strategies, we designed a phototreatment panel that covered chemical and mechanistic diversity. We therefore selected five complementary photodynamic approaches applied under both short-term (30 min) and prolonged (7–8 h) exposure conditions. This combination enabled us to capture both acute stress responses and longer-term adaptive trajectories across fundamentally different modes of photodynamic action.

Three of the selected treatments employed exogenous photosensitizers representing distinct and well-characterized chemical classes with different photochemical properties. TMPyP [5,10,15,20-tetrakis(1-methyl-4-pyridinium)porphyrin tetra-(p-toluenesulfonate)] is a tetra-cationic porphyrin derivative that acts predominantly via a Type II photodynamic mechanism through high-efficiency singlet oxygen generation [7]. Rose Bengal, a halogenated fluorescein derivative with strong absorption in the green spectral range (480–550 nm), also primarily drives Type II photochemistry but differs markedly from TMPyP in molecular structure and charge distribution [7]. In contrast, new methylene blue (NMB), a phenothiazinium dye absorbing in the red region, operates mainly via a Type I mechanism involving electron transfer and radical formation [15]. Together, these three photosensitizers span the major classes of clinically and experimentally relevant photodynamic agents and encompass both dominant photochemical reaction pathways. To complement these exogenous approaches, we included two strategies based on endogenous bacterial chromophores. Antimicrobial blue light (aBL) exploits naturally occurring intracellular porphyrins, providing a photosensitizer-free modality that is increasingly explored for clinical and environmental applications. In parallel, we combined aBL with 5-aminolevulinic acid (ALA) to enhance intracellular accumulation of endogenously produced porphyrins, thereby intensifying photodynamic effects while preserving the same light-based activation principle [16].

To our knowledge, this is the first study to systematically characterize the transcriptomic response of *E. coli* to photodynamic strategies. This study provides a comprehensive, time-resolved transcriptomic framework for understanding how *E. coli* responds to different photodynamic antimicrobial treatments that rely on both exogenous and endogenous photosensitization pathways. By comparing short- and long-term exposures across mechanistically distinct phototreatments, we identify common stress responses as well as treatment-specific adaptive changes that influence bacterial survival under photodynamic stress. These findings improve our fundamental understanding of light-based antimicrobial action and support the rational development and standardization of aPDI and aBL for translational and clinical applications

## Materials and Methods

### Strain and culture conditions

*E. coli* BW25113 Keio collection parent strain was used [17]. Bacteria were cultured in M9 minimal medium supplemented with 0.4% glucose, 0.2 mM MgSO_4,_ and 0.1 mM CaCl_2_ or seeded on M9 agar plates with the same supplements as in liquid medium and 1.5% agar. For overnight culture, a single colony was inoculated into 5 ml of M9 or LB medium and incubated at 37°C for 18 hours under aerobic conditions with shaking (150 rpm). Log-phase culture was prepared by diluting the overnight culture 1:25 into 10 mL of fresh supplemented M9 medium and incubating for approximately 5.5 h under the abovementioned conditions.

### Light sources

Three custom-made light-emitting diode (LED) lamps were used in this study with emission maxima at 415 nm and a radiosity of 25 mW/cm^2^ (Cezos, Poland), λ_max_ 522 nm and a radiosity of 10.6 mW/cm^2^ (Cezos, Poland), and λ_max_ 633 nm and a radiosity of 138.3 mW/cm^2^ (lab255, Poland). A 415 nm lamp was used for aBL and aBL with ALA treatment, while a 522 nm lamp was used for RB, and a 633 nm lamp was used for TMPyP and NMB treatment.

### Chemical compounds

All compounds were purchased from Merck (Sigma-Aldrich, Darmstadt, Germany). Stock solutions were prepared in sterile water and stored in the dark at 4°C - 1 mM for photosensitizers and 5 mg/ml for ALA. Working solutions were freshly prepared by diluting the stock solutions to the desired concentrations.

### Determination of the sub-lethal conditions of photodynamic treatments for short- and long-term duration

For the short-term treatment, overnight cultures were grown in LB medium for the aBL group, and in supplemented M9 medium for aPDI and aBL + ALA groups. Then, cultures were diluted 1:25 into 10 mL of fresh, supplemented M9 medium, and 5 mL aliquots were transferred into a 6-well plate. At this point, for aBL-ALA treatment, cultures were supplemented with the desired concentration of ALA. The plate was then placed in a ThermoMixer® C (Eppendorf, Germany) at 37 °C for approximately 5.5 h with continuous shaking and heating. After incubation, the OD_600_ was measured, and cultures were adjusted to an optical density of 0.3 in M9 medium, and aliquots of 900 μl were transferred to a 24-well plate. For aBL/aBL + ALA samples, cultures were immediately subjected to illumination, while for aPDI samples, the PS was added to the bacterial suspensions at the desired concentration, followed by a 15-minute incubation in the dark at 37 °C prior to illumination. After irradiation, 10 μl aliquots were serially diluted tenfold in PBS to generate dilutions of 10^−1^ to 10^−5^ and streaked horizontally onto M9 agar plates. Plates were incubated at 37 °C for 16–20 h, after which colony-forming units (CFUs) were counted to determine survival rates. Control groups included cells that were not treated with PSs or light. Each experiment was performed in triplicate.

For the long-term treatment, overnight cultures were grown in different media depending on the treatment group: in LB medium for the aBL group, in supplemented M9 medium with the addition of ALA for the aBL+ALA group, and in supplemented M9 medium for the aPDI groups. Then cultures were similarly diluted 1:25 into 10 mL of fresh supplemented M9 medium, and 900 μl aliquots were transferred to a 24-well plate. For the aBL and aBL + ALA groups, the plate was immediately placed in the ThermoMixer® C and subjected to simultaneous incubation and illumination. In the aPDI groups, photosensitizers were added at the desired concentrations, followed by 15 minutes of dark incubation at 37 °C, after which the samples were illuminated and incubated simultaneously. Optical density (OD_600_) was measured every hour using a microplate reader (Envision, PerkinElmer) to monitor bacterial growth for 12 h. Control groups included cells that were not treated with PSs or light. Each experiment was performed in triplicate. The detailed experimental setup is presented in Figure 1.

**Figure 1.**
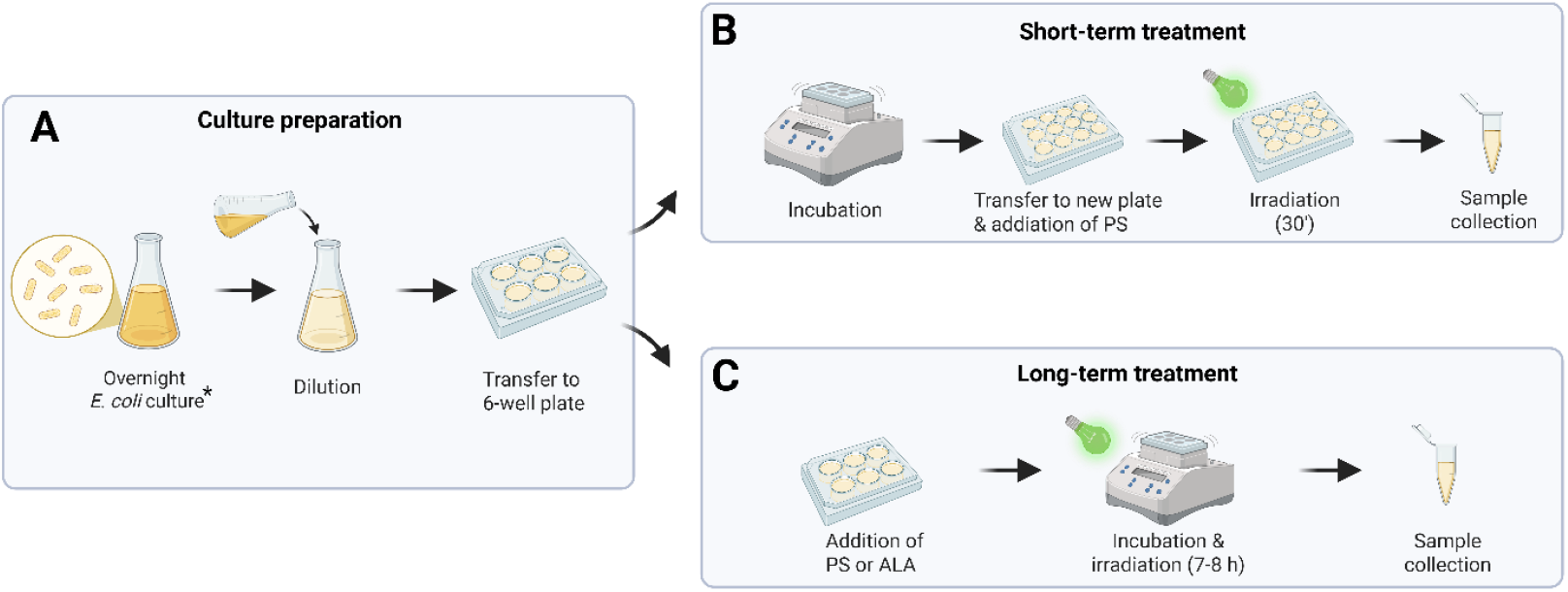
Experimental setup for photodynamic treatment.

### RNA extraction and sequencing

Total RNA was isolated from *E. coli* cells after illumination using the RNeasy^®^ Mini Kit (Qiagen, Germany). For samples exposed to short-term photodynamic treatment, an additional 40-minute incubation at 37 °C post-irradiation was included to allow for cellular recovery. RNA concentration was measured using a NanoDrop One spectrophotometer (Thermo Scientific, USA), and samples were stored at −80 °C until further processing. RNA samples were sent to Lexogen GmbH (Austria), where quality control, genomic DNA removal, rRNA depletion (RiboCop), and library preparation were performed using the CORALL Total RNA-Seq Library Prep Kit. Sequencing was conducted on an Illumina Novaseq X platform in paired-end 150 bp read mode, yielding an average of 10 million reads per sample.

### Transcriptomic data analysis

The RNA-seq reads were subjected to quality control using FastQC v0.12.1. Subsequently, quality filtering and adapter trimming were performed with BBDUK2 from BBMAP package v36.14, using the following parameters: qtrim = w, trimq = 20, maq = 10, forcetrimleft=20, k = 23, mink = 8, hdist = 1, tbo, tpe, minlength = 100, removeifeitherbad = t. High-quality reads were then mapped to the complete sequenced genome of the reference strain K-12 (ENSEMBL ASM584v2) using Bowtie 2. The filtered FASTQ files were used as input for read counting using RSEM v1.2.30. Differential expression analysis was performed using DESeq2 v1.34.0, and genes with a > 1.5-fold change in expression and an adjusted P-value < 0.05 were considered as differentially expressed (DE). Further, functional analysis of enrichment in Gene Ontology (GO) terms and Kyoto Encyclopedia of Genes and Genomes (KEGG) pathways was conducted with clusterProfiler, KEGGREST, and org.EcK12.eg.db libraries in Bioconductor/R.

Specifically, enrichment was assessed across Biological Process (BP), Molecular Function (MF), and Cellular Component (CC) domains using Benjamini-Hochberg (BH) adjusted p-values and q-values with a cutoff of 0.05. To ensure accurate database cross-referencing, gene symbols were converted to Entrez IDs using the *bitr* utility. Significantly enriched terms were visualized as dot plots (clusterProfiler), prioritizing the top 25 categories for each functional domain.

Regulon enrichment analysis was performed to identify transcription factors (TFs) significantly associated with differentially expressed genes (DEGs) using regulatory interaction data from RegulonDB. DEGs were filtered based on an adjusted p-value < 0.05 and ∣log2Fold Change∣ > 1. Enrichment was calculated using a Fisher’s exact test, with p-values corrected for multiple testing using the Benjamini-Hochberg procedure. The analysis was restricted to TFs with at least 5 target genes in the background set.

## Results

### Optimization of sub-lethal photodynamic treatment parameters for *E. coli* transcriptomic profiling

To obtain biologically meaningful transcriptomic data, it was essential to first establish sub-lethal photodynamic conditions that elicit cellular stress responses without causing extensive cell death or lysis. Careful optimization of these parameters ensures that observed gene expression changes reflect active bacterial adaptation rather than nonspecific damage or loss of viable cells. This step is therefore critical for accurately interpreting the molecular mechanisms underlying bacterial responses to photodynamic treatments.

For short-term treatments, sub-lethal doses were defined as those that resulted in an approximately 1 log_10_ reduction in CFU/ml in the mid-log phase of growth during 30 minutes of exposure. The duration of the treatment was selected based, among others, on previous study reports that 15-30 minutes is sufficient for activation of stable gene expression, including initiation of SOS response, and enables monitoring of rapid bacterial response. Moreover, the threshold of 1 log_10_ CFU/ml in viability allows for an accurate mechanistic analysis without being biased by irreversible cell damage or lysis. In terms of long-term treatments, sub-lethal doses were defined as continuous exposure until cultures reached the mid-exponential phase of growth, corresponding to approximately 50% inhibition of growth (based on OD_600_ values) compared to untreated controls. Control cultures were grown for 6 h, while treated cultures required 7-8 h to reach the same phase. The duration of treatment was chosen based on observed bacterial growth dynamics, ensuring that cells reached the log phase, which is optimal for high RNA yield from metabolically active cells. Additionally, the extended exposure of low-dose aPDI/aBL allows for the assessment of long-term or adaptive responses that may develop over several bacterial generations. The estimated sub-lethal conditions for the abovementioned treatments are presented in Table 1.

**Table 1.**
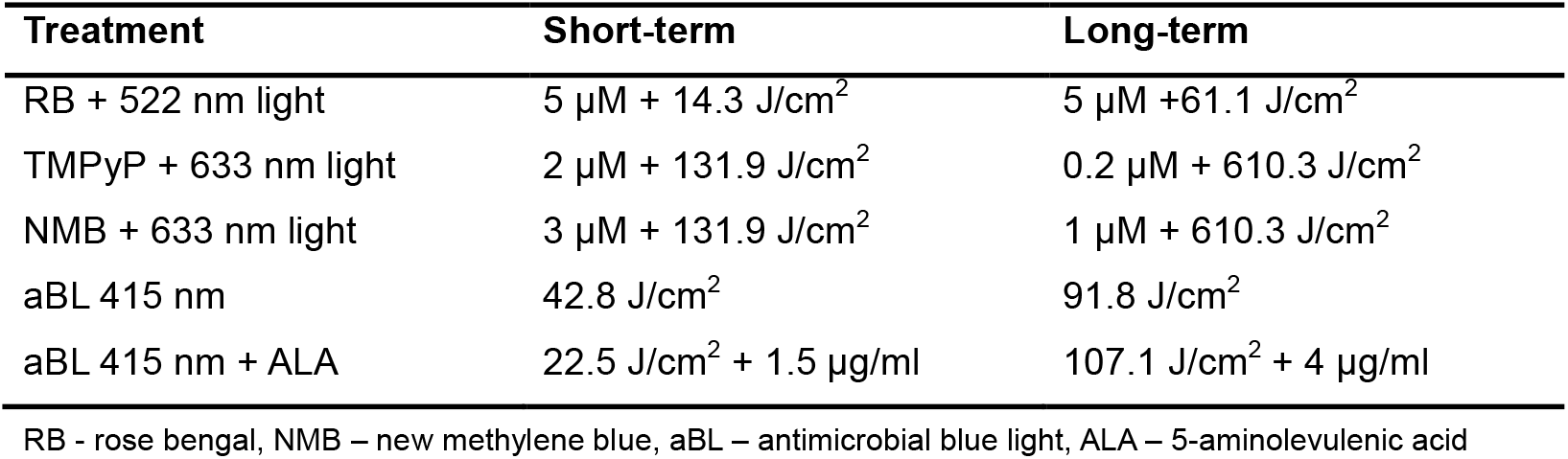
Experimental conditions for short- and long-term photodynamic treatments.

### Exposure duration determines the strength and specificity of *E. coli* transcriptomic responses

To obtain a comprehensive picture of how *E. coli* responds at the transcriptomic level to various photodynamic protocols, RNA-seq sequencing was performed. The detailed experimental setup is presented in Figure 1. RNA-seq data were obtained for all conditions with high coverage and mapping quality, and detailed sequencing statistics are provided in the Supplementary Materials Table S1 and Figure S1, S2.

Principal component analysis (PCA) of log_2_-transformed gene expression values from control and treated samples (short exposure conditions, Figure 2A) revealed that the five treatment groups formed distinct clusters, with biological replicates tending to group within those clusters, suggesting a strong transcriptional response to those conditions. Notably, the treatment clusters also showed a directional tendency away from the control. In the case of long treatment (Figure 2B), the differentiation between control and treated samples is less pronounced, but we observed a strong separation of aBL-treated samples from the other groups. This suggests that prolonged photodynamic exposure exerts a weaker effect on gene expression, except for blue light. As a complement to the PCA analysis, a hierarchical clustering heatmap of the top 100 most variable genes (selected based on variance of mean expression values across three biological replicates per group) revealed a clear separation between control and treated samples following short exposures (Figure 2C), with distinct gene expression profiles induced by treatment. However, samples did not cluster according to the type of treatment (aBL/aBL+ALA vs. aPDI). In long-treated samples (Figure 2D), clustering indicated a less robust transcriptome shift for extended exposure of aBL+ALA and aPDI. Notably, cells subjected to blue light showed the most distinct transcriptional profile among all samples. *E. coli* cells exposed to short-duration aBL or aPDI treatments exhibited extensive transcriptomic change (Figure 2E, Table 2), with an average of 2,105 differentially expressed genes (padj < 0.05; fold change ≥ 1.5), accounting for 49.1% of all actively transcribed genes. The strongest response was observed for TMPyP combined with red light (57.9%), while RB activated by green light elicited the weakest effect (34.9%). In contrast, for all long treatments, we observed markedly reduced response in the number and proportion of DEGs, with an average of only 16.9% of the transcriptome being differentially expressed. The weakest response was marked in the extended TMPyP treatment, affecting just 6.6% of the transcriptome. It is worth noting that despite the variability in DEG numbers across conditions, the ratio of upregulated to downregulated genes remained relatively balanced in most treatments.

**Table 2.** Differential gene expression profiles under phototreatments.

| Treatment | No. of upregulated genes | No. of downregulated genes | No. of differentially expressed genes (% of transcriptome) |
| --- | --- | --- | --- |
| aBL [S] | 1210 | 1223 | 2433 (56.7%) |
| aBL [L] | 415 | 407 | 822 (19.2%) |
| aBL + ALA [S] | 1130 | 1075 | 2205 (51.4%) |
| aBL + ALA [L] | 332 | 269 | 601 (13.9%) |
| RB [S] | 829 | 667 | 1496 (34.9%) |
| RB [L] | 543 | 494 | 1037 (24.1%) |
| NMB [S] | 1021 | 948 | 1969 (46.1%) |
| NMB [L] | 193 | 92 | 285 (6.6%) |
| TMPyP [S] | 1253 | 1173 | 2426 (57.9%) |
| TMPyP [L] | 540 | 349 | 889 (20.7%) |

S- short treatment, L-long treatment
% of transcriptome was calculated as % number of DEGS relative to number of active genes in the transcriptome (baseMean > 0)

**Figure 2.**
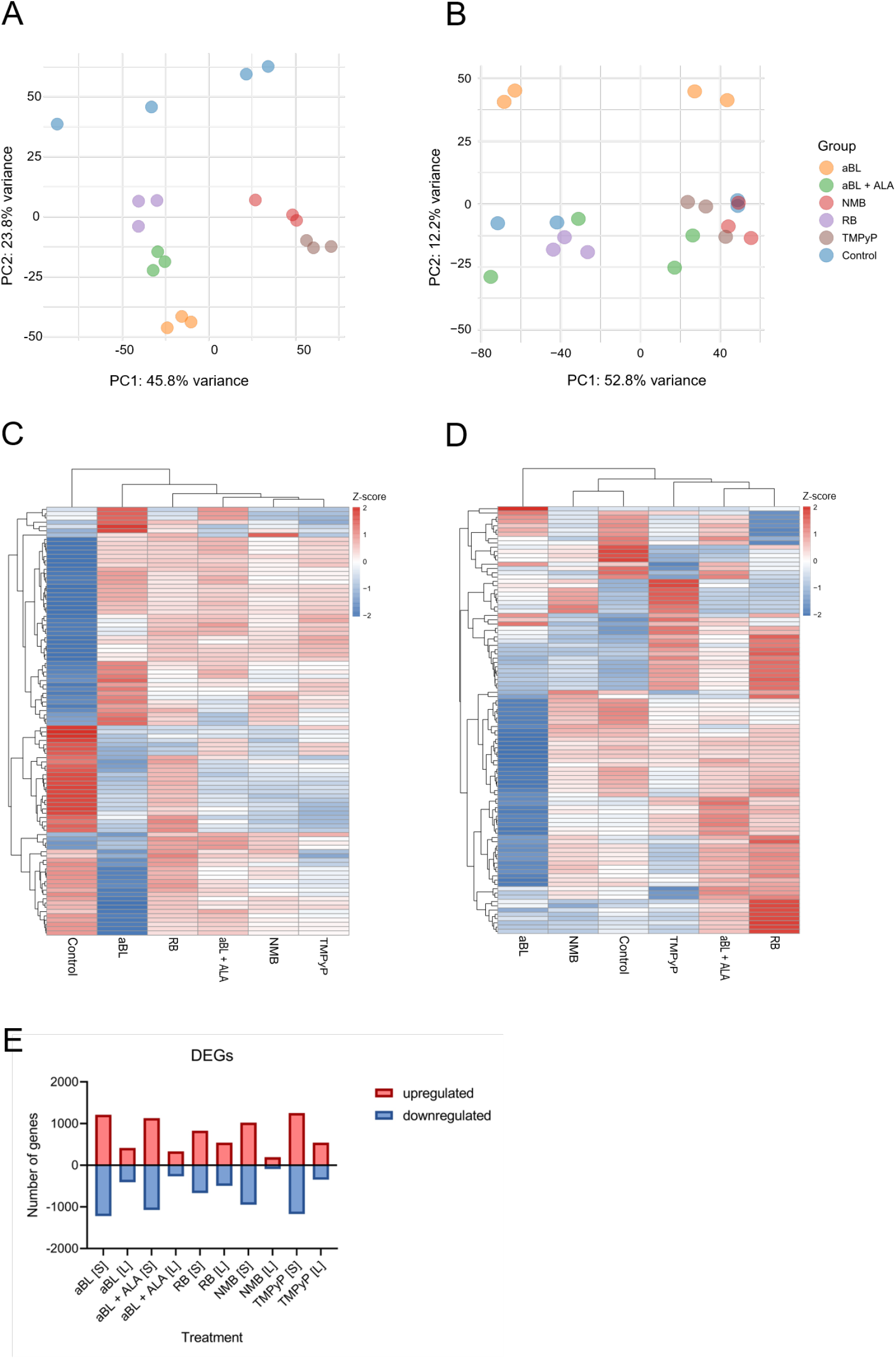
Transcriptional response of *E. coli* BW25113 to ten photodynamic treatments (short and long exposures – RB + green light, TMPyP and NMB + red light, aBL and aBL + ALA. **(A–B)** Principal component analysis (PCA) of normalized gene expression data for short (A) and long (B) treatment conditions. Each point represents a biological replicate. **(C-D)** Heatmaps of the top 100 most variable genes across experimental groups, based on variance of mean expression values (averaged across three biological replicates per group). Gene-wise expression values were standardized (z-scores), and both genes and groups were hierarchically clustered using Euclidean distance, across short-(C) and long-term (D) treatment, respectively. **(E)** Differentially expressed genes (DEGs) per treatment. Bars show the total number of DEGs (padj < 0.05; fold change ≥ 1.5), with red and blue indicating upregulated and downregulated genes, respectively. Results shown are based on at least three independent biological replicates.

### Short-term photodynamic stress triggers rapid defense pathways, while prolonged exposure drives metabolic adaptation

To investigate the functional impact of various photodynamic treatments on *E. coli* cells, gene ontology (GO) enrichment analysis and KEGG pathway analyses were performed on differentially expressed genes (DEGs) identified for all short- and long-term conditions (Figures 3–4).

**Figure 3.**
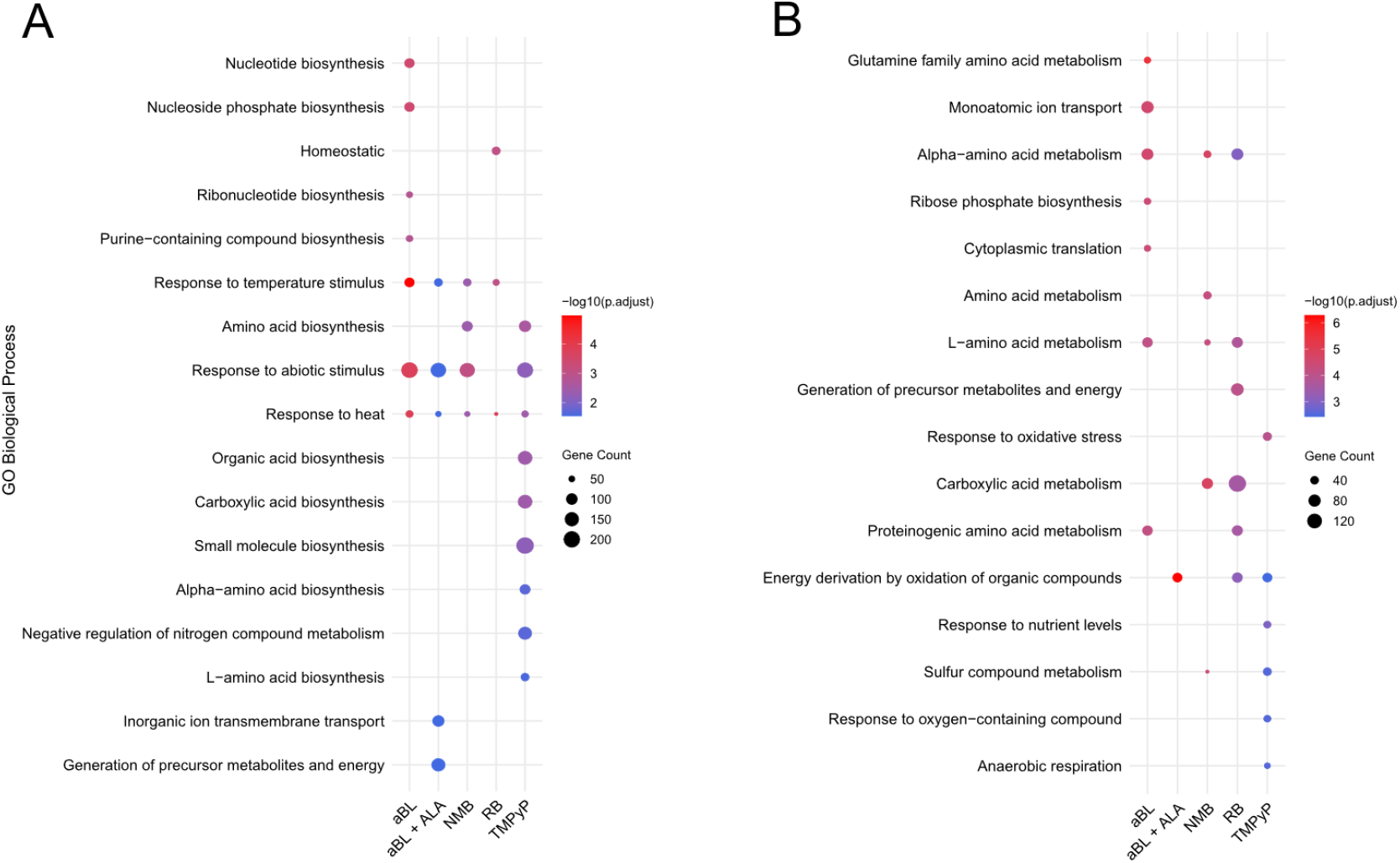
Gene Ontology (GO) enrichment analysis of differentially expressed genes (DEGs) in *E. coli* BW25113 following short- and long-term photodynamic treatments. **(A)** GO terms significantly enriched among DEGs after short-term exposures (30 min) to sub-lethal photodynamic stress. **(B)** GO terms significantly enriched among DEGs after long-term exposures (8-9 h). Dot size represents the number of DEGs annotated to the term, and color indicates the adjusted *p*-value (padj). Results shown are based on at least three independent biological replicates.

GO analysis revealed that short-term exposures predominantly affected genes involved in the biosynthesis of core macromolecular precursors, including amino acids, nucleotide bases, and intermediary metabolites (Figure 3A). Notably, processes linked with response to temperature stimulus, heat, and abiotic stress were significantly overrepresented for most of the treatments, indicating rapid activation of general stress response pathways. Prolonged exposures instead showed significant overrepresentation of pathways associated with metabolic remodeling across nearly all experimental conditions (Figure 3B). Observed enrichment in GO terms related to organic acid metabolism, energy derivation by oxidation of organic compounds, and generation of precursor metabolites and energy suggests a pronounced shift in cellular metabolism across all conditions as a prolonged adaptation to stress. The comparative analysis of short-vs long-term responses suggests that bacterial cells after temporary treatment prioritize immediate survival, while prolonged exposure leads to an altered metabolic state.

While GO enrichment analysis was performed on all differentially expressed genes to capture the overall biological processes affected by phototreatments, KEGG pathway enrichment was conducted separately for up-(Figure 4A and 4C) and down-regulated (Figure 4B and 4D) genes to preserve the directionality of transcriptional responses at the pathway level. In short-term treatments (Figure 4A), up-regulated genes showed consistent overrepresentation of pathways related to exopolysaccharide biosynthesis, sulfur metabolism, and two-component regulatory systems. These pathways are associated with canonical stress response, including biofilm formation and environmental sensing, as well as sulfur metabolism can represent involvement in oxidative stress and detoxification. Notably, RB-mediated aPDI was associated with the broadest functional diversity among upregulated pathways, including unique activation of fatty acid degradation, lysine degradation, and methane, nitrogen, and propanoate metabolism, suggesting induction of alternative energy and detoxification routes. In contrast, NMB-mediated aPDI and aBL combined with ALA showed enrichment of fewer distinct pathways, which may reflect a more focused or limited transcriptional rearrangement. Downregulated genes in short-term exposures (Figure 4B) exhibited consistent enrichment in ABC transporters, biosynthesis of cofactors, and secondary metabolites biosynthesis across all five treatment conditions. Additionally, most phototreatments showed suppression of amino acid biosynthesis, carbon fixation, and oxidative phosphorylation. These findings suggest that photooxidative stress triggers in *E. coli* cells global metabolic downshift, likely reflecting a transition from growth to survival state. Interestingly, RB-mediated aPDI, which exhibited the strongest transcriptional activation, showed the lowest number of down-regulated pathways, whereas the remaining treatments exhibited a wider range of suppressed processes.

**Figure 4.**
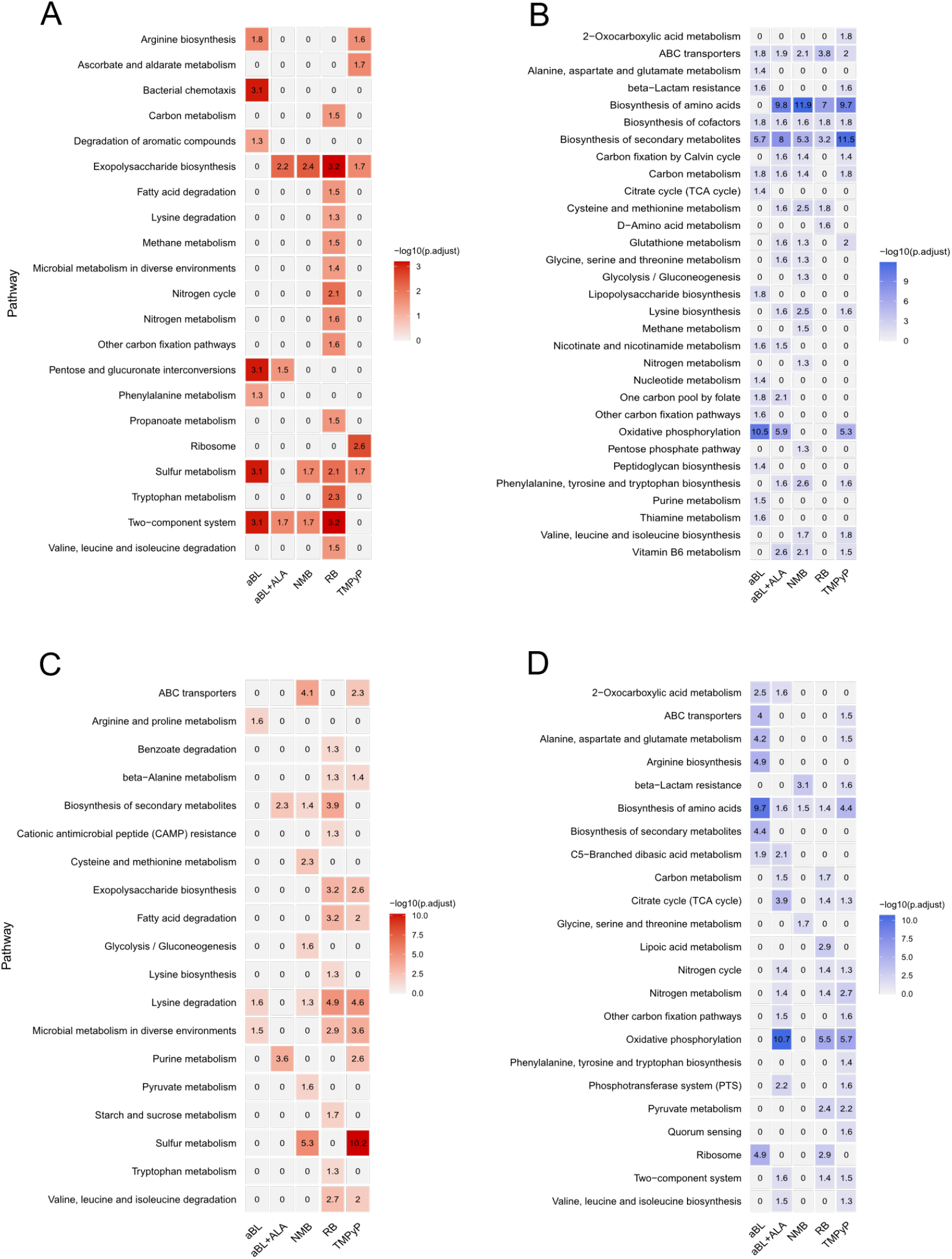
KEGG pathway enrichment of differentially expressed genes in *E. coli* BW25113 following photodynamic treatments. Pathway enrichment was analyzed separately for upregulated (A, C) and downregulated (B, D) genes. **(A–B)** Short-term treatments. **(C–D)** Long-term treatments. Results shown are based on at least three independent biological replicates.

Long-term exposure to the five photodynamic conditions resulted in a distinct transcriptional profile compared to short-term responses. Among up-regulated genes (Figure 4C), lysine degradation was the most consistently enriched pathway, indicating a shift toward amino acid catabolism. Additionally, secondary metabolite biosynthesis and microbial metabolism in diverse environments were enriched in 3 out of 5 conditions, reflecting enhanced metabolic flexibility and environmental adaptation. The blue light irradiation with or without ALA exhibited the smallest set of enriched pathways, while all aPDI treatments triggered more extensive gene activation, reflected by a broader range of enriched pathways. Down-regulated genes under long-term treatment (Figure 4D) showed a pattern partially similar to the short-term response. The biosynthesis of amino acids was consistently downregulated across all conditions, indicating a suppression of growth-related anabolic processes. Moreover, we observed frequent downregulation of central metabolic pathways, including the TCA cycle, oxidative phosphorylation, nitrogen cycle, and two-component systems, suggesting a deep metabolic reprogramming characterized by reduced respiration, nitrogen turnover, and signal transduction activity. All prolonged phototreatments, except NMB-mediated aPDI, manifested a similar level of pathway enrichment.

Notably, a consistent pattern between GO and KEGG pathway enrichment analyses was observed. Both approaches revealed that short-term treatments were associated with the activation of stress-related processes such as response to heat and response to abiotic stimulus (Figure 3A and Figure S3A), as well as sulfur metabolism, two-component system, and exopolysaccharide biosynthesis (Figure 4A). These responses were accompanied by the suppression of biosynthetic and energy-consuming processes, including nucleoside phosphate metabolism, and small molecule biosynthesis (Figure S3B), as well as amino acid biosynthesis, carbon metabolism, and ABC transporters (Figure 4B). In long-term treatments, both analyses supported a shift toward metabolic reprogramming and utilization of alternative energy sources. For instance, GO terms related to ribosome biogenesis, translation, and respiration were significantly downregulated (Figure S3D), aligning with the suppression of KEGG pathways such as ribosome, oxidative phosphorylation, and the TCA cycle (Figure 4D), while upregulated genes showed enrichment in lysine degradation and fatty acid metabolism (Figure 4C).

### Distinct regulatory programs underlie short- and long-term responses to photodynamic stress

To further explore the regulatory mechanisms underlying the observed transcriptional changes, we performed transcription factor regulon enrichment analysis across all short- and long-term photodynamic treatments (Figure 5). Under short exposure conditions, regulon enrichment was predominantly focused on regulators associated with acid stress, metal homeostasis, and envelope integrity. Across most treatments, GadX, GadW, and RcsAB emerged as the most consistently enriched regulons. In parallel, multiple regulators associated with envelope and membrane stress responses, most notably RcsAB, CpxR, and BaeR, were detected. Short-term exposure also elicited strong enrichment of global metabolic and redox regulators, including Fur and Cbl. Notably, relatively narrow regulon engagement was observed, with minimal involvement of global transcriptional regulators, suggesting that early responses primarily rely on specialized stress-response modules.

**Figure 5.**
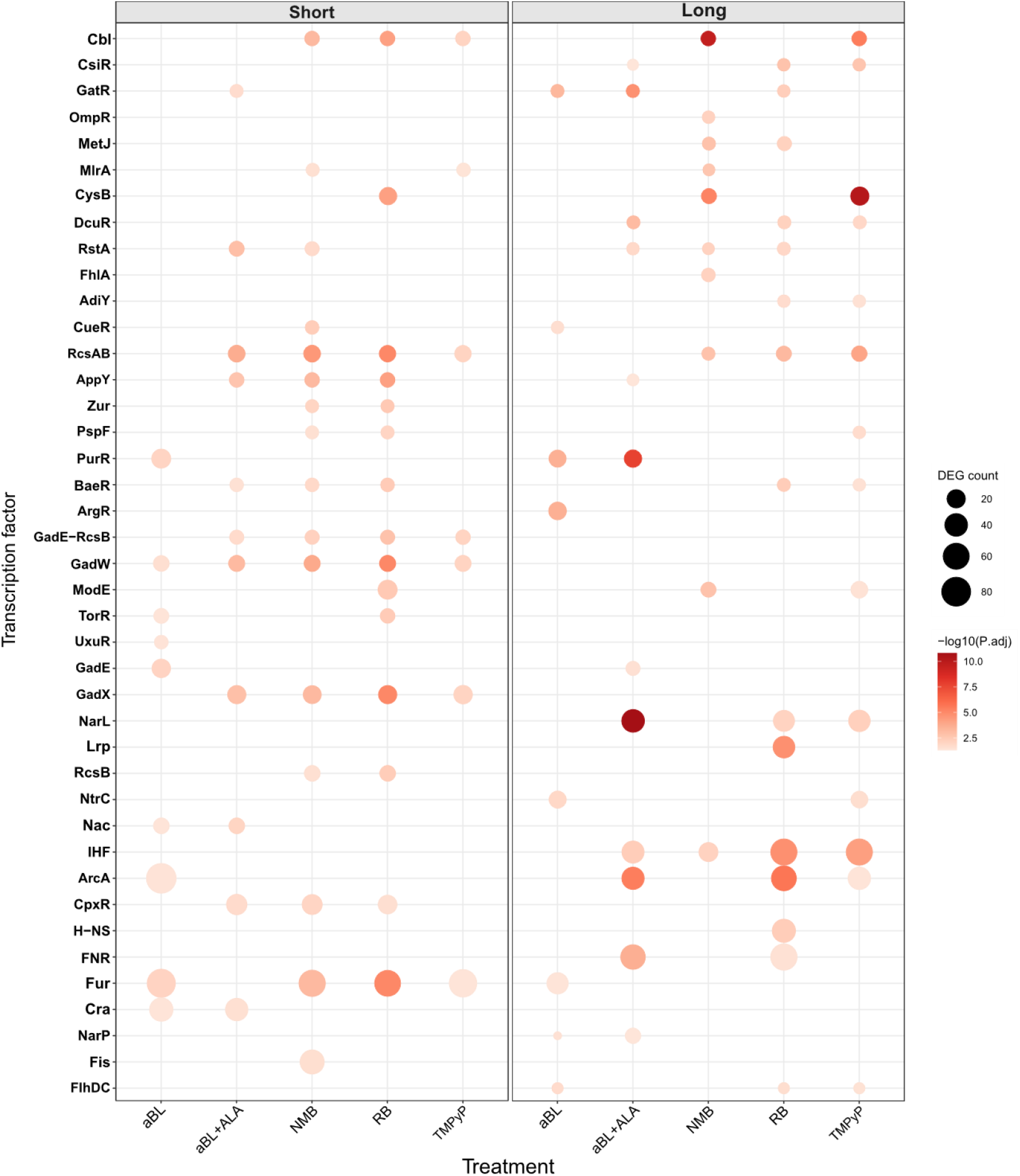
Enrichment of transcription factor regulons under short- and long-term photodynamic treatments. Dot size represents DEG count, and color indicates –log10 adjusted P-value.

In contrast, long-term treatment induced enrichment of global regulators associated with central metabolism, anaerobic respiration, and nitrogen utilization. Across multiple treatments, ArcA, IHF, FNR, and NarL emerged as dominant regulons, indicating a shift toward systemic transcriptional reprogramming. The diversity of regulatory responses under prolonged exposure was exemplified by treatment-specific signals, including the enrichment of OmpR and FhlA under NMB conditions, and of Fur under aBL conditions.

Together, these results demonstrate that exposure duration is a primary determinant of transcriptional regulatory architecture. Short-term responses rely on specialized stress regulons, whereas long-term adaptation involves extensive engagement of global metabolic and respiratory regulators. Functional classification of enriched regulons is provided in Table S2.

### Short-term convergence and long-term divergence in photodynamic responses

To evaluate the overlap and divergence in transcriptional responses to the five treatments, we constructed Venn diagrams of differentially expressed genes for short-term (Figure 6A) and long-term (Figure 6B) exposures. Short-term treatments revealed a substantial common response, with 891 shared DEGs across all conditions, indicating a robust common core response. Among treatment-specific profiles, aBL induced the largest set (403 genes), followed by TMPyP (197), aBL+ALA (99), NMB (40), and RB (35). In contrast, long-term exposures showed markedly reduced overlap, with only 46 shared DEGs. Unique responses were more pronounced, particularly for aBL (350), RB (262), and TMPyP (159), whereas aBL+ALA (103) and NMB (15) exhibited smaller exclusive sets. These patterns suggest that the extensive overlap in shorter treatments reflects conserved stress-responsive mechanisms, while the divergence during prolonged treatment highlights distinct adaptive responses to each phototreatment.

**Figure 6.**
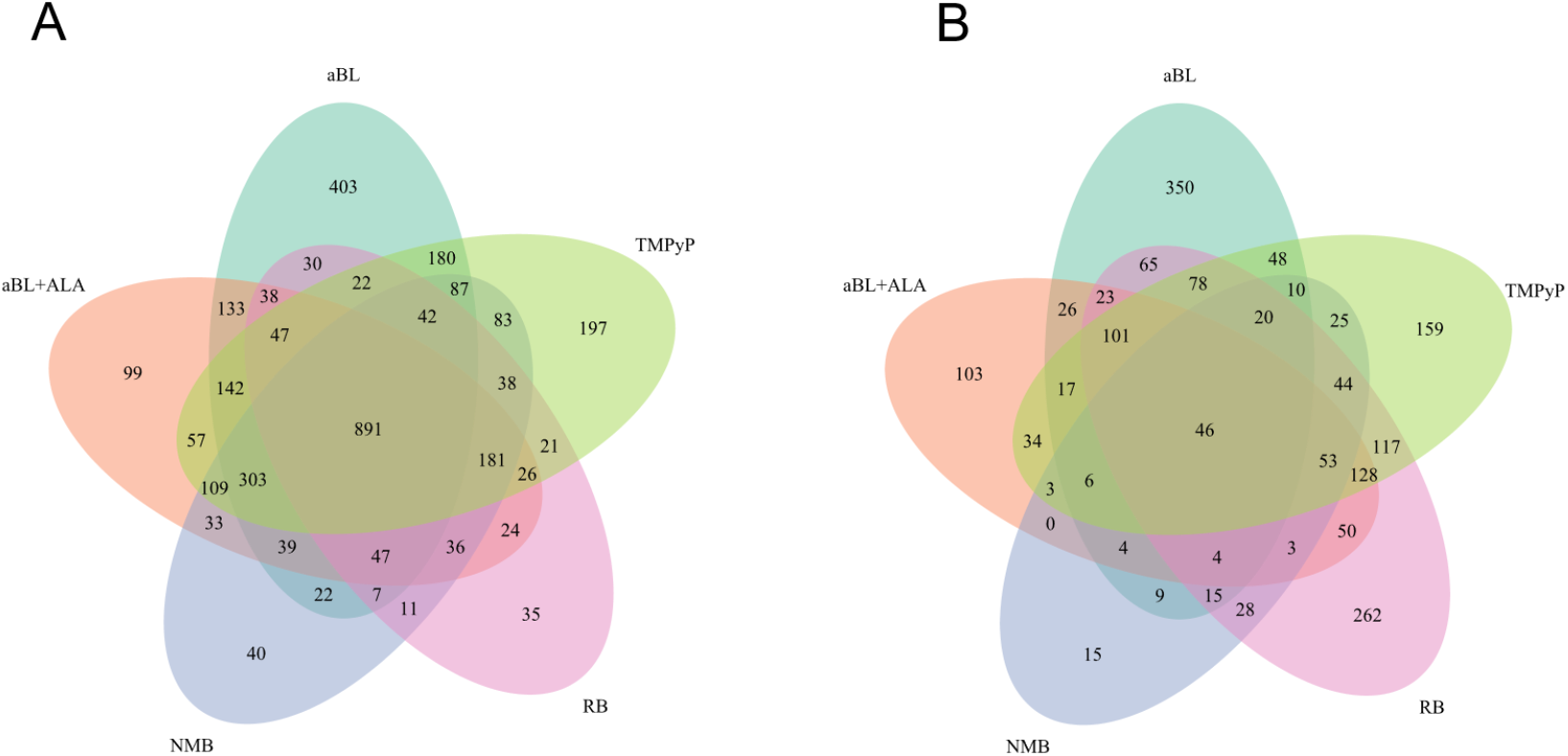
Venn diagrams showing the overlap of differentially expressed genes (DEGs) across five phototreatment conditions under (A) short-term and (B) long-term exposure. DEGs were defined as those with an adjusted pvalue (padj) < 0.05 and a fold change ≥ 1.5. Results are based on a minimum three independent biological replicates.

## Discussion

Specifically, the aim of this study was to characterize the transcriptome response of *Escherichia coli* to photodynamic inactivation during both short-term and long-term exposure. Analysis of these different treatment conditions is crucial, as microorganisms exhibit different transcriptional responses depending on the duration of the stress factor. The use of a single time point can therefore lead to incomplete or distorted conclusions [18].

Sublethal treatment conditions have been carefully defined to enable accurate and correct interpretation of transcriptome data and were consistent with other transcriptome studies on bacteria exposed to sublethal photodynamic or chemical stress [11], [18]. Ultimately, the RNA-seq data obtained confirmed the validity of the experimental conditions, as short-term treatment elicited a broad stress response in the microorganism affecting approximately 35– 58% of the transcriptome, which is characteristic of acute stress exposure, while long-term exposure triggered a weaker but distinct adaptive response, including metabolic restructuring. Such results would not have been possible if the treatment conditions had represented lethal exposure. The above arguments demonstrate that the sublethal conditions were correctly identified.

A global transcriptome analysis showed that *E. coli* undergoes an extensive and exposure-duration-dependent response to photodynamic stress. Short-term exposure triggered transcriptional changes affecting up to 58% of the genes, while long-term exposure triggered a smaller number of transcriptional changes, but with a more pronounced adaptive character. This transition from an acute to an adaptive response reflects observations described for other bacteria exposed to oxidative or photooxidative stress [13], [14], [19].

The comparative transcriptome analysis revealed both common and treatment-specific responses of the bacteria to photodynamic inactivation (aPDI) and antibacterial blue light (aBL). Short-term exposure triggered a relatively similar stress program under all conditions studied, while long-term exposure revealed different adaptation strategies. Short-term treatment showed a preserved stress program characterized by the activation of sulfur metabolism, exopolysaccharide biosynthesis, and two-component regulatory systems, accompanied by an inhibition of anabolic processes such as amino acid biosynthesis and oxidative phosphorylation (Figure 4AB). These changes indicate a rapid slowing of metabolism and the activation of defense mechanisms, which is consistent with previous studies on the response to blue light-induced stress conditions in *Staphylococcus aureus* [12], *E. coli* [20], *Campylobacter jejuni* [14] and *Acinetobacter baumannii* [21]. In contrast, longer exposure showed a more diverse response with unique transcriptional signatures for each treatment (Figure 6B). In particular, long-term exposure to aBL and RB led to transcriptomic changes in the largest group of treatment-specific genes, suggesting strong adaptive diversification. This likely reflects the utilization of universal defense mechanisms and the emergence of specialized survival strategies.

One of the most striking observations is the transcriptional response triggered by aBL. Transcriptome analysis showed that the response of *E. coli* to aBL varies considerably depending on the duration of exposure. Short-term exposure leads to a strong downregulation of genes for oxidative phosphorylation but has no effect on amino acid biosynthesis (Figure 4B), while long-term exposure has the exact opposite effect, i.e., a strong suppression of amino acid biosynthesis without significant changes in oxidative phosphorylation (Figure 4D). These opposing effects suggest a time-dependent metabolic reprogramming under the influence of photodynamically induced stress. During short-term exposure to aBL, light-induced endogenous chromophores (e.g., flavins, porphyrins) generate reactive oxygen species (ROS) and trigger an acute response to oxidative stress. The observed inhibition of oxidative phosphorylation likely reflects a protective mechanism that limits ROS production from the electron transport chain, while biosynthetic pathways remain active to support protein synthesis and repair. However, long-term exposure to aBL leads to an accumulation of oxidative damage that impairs biosynthesis capacity. A substantial reduction in amino acid biosynthesis suggests a transition to a state of metabolic inactivity or dormancy, in which microorganisms conserve resources and prioritize basic functions. Stable expression of oxidative phosphorylation genes during this phase may represent an adaptation to maintain minimal energy production. Together, these results confirm a two-phase adaptation model: an early phase of redox defense characterized by temporary inhibition of respiration, followed by a late phase of resource conservation characterized by inhibition of anabolic metabolism and stabilization of energy pathways. A comparable light-dependent disruption of electron transport was also observed in *C. jejuni* exposed to violet-blue light, with inactivation of enzymes containing iron-sulfur clusters and loss of heme from cytochromes, leading to disruption of energy metabolism [14]. Similarly, under the influence of blue light, *S. aureus* showed transcriptional changes that led to a reduction in the activity of growth pathways [12], [13].

Another unique response was observed during photodynamic inactivation with TMPyP. In this case, activation of sulfur metabolism was observed (Figure 4C). The upregulation of sulfur pathways likely reflects an increased need for thiol-based redox buffering, consistent with the role of glutathione and cysteine metabolism in detoxifying reactive oxygen species [19]. These different transcriptomic response patterns underscore that while all treatments studied exert photooxidative pressure, the specific metabolic adaptations depend on the photosensitizer used and the corresponding wavelength of light.

Exposure of microorganisms to various stress factors has shown that, despite stress factor-specific transcriptome responses, there is a transcription response common to the various factors that involves the upregulation of genes involved in stress protection, repair, and detoxification, as well as the downregulation of genes associated with rapid growth, protein synthesis, and energy-intensive biosynthesis pathways [22]. Heat shock is one of the best-characterized environmental stressors at the transcriptome level. In bacteria such as *E. coli*, a sudden increase in temperature rapidly induces the alternative sigma factor σ32, which activates the transcription of classic heat shock genes, including *dnaK, groEL, groES*, and *clpB* [23]. These encode chaperone molecules and proteases required for the prevention of protein aggregation and the refolding of denatured proteins. Transcriptomic analyses also show repression of ribosomal protein genes and translation-related genes, which, similar to our findings, reflects the transition from growth metabolism to adaptive metabolism [22]. Next, oxidative stress caused by reactive oxygen species (ROS), also triggers a significant transcriptional response. In *E. coli*, this leads to the activation of the OxyR and SoxRS regulons, resulting in strong induction of antioxidant and redox-balancing genes, including *katG* (catalase), *ahpC* (alkyl peroxide reductase), *sodA*, and *sodB* (superoxide dismutase) [24]. Another stress agent, i.e., exposure to ultraviolet (UV) radiation and other DNA-damaging factors triggers transcriptome activation of DNA repair and tolerance pathways, which are mainly dependent on the SOS response [25], [26]. At the same time, genes associated with cell cycle progression and replication are suppressed, reflecting a transcriptional strategy that prioritizes genome stability over proliferation [22]. All these are in line with our analyses regarding transcriptomic response over time, revealing that the responses of microorganisms are highly dynamic and characterized by rapid induction of early regulatory and signaling genes, followed by a slower reorganization of metabolism.

Understanding which regulatory networks are activated under different photodynamic exposure regimes is essential for deciphering how bacteria sense and ultimately survive light-induced stress. During short-term photodynamic exposure, enrichment analysis mainly identified stress-response regulons linked to acid stress (GadW, GadX), envelope integrity (RcsAB, CpxR, BaeR, PspF), and metal or redox homeostasis (Fur, Cbl) (Figure 5). This regulatory pattern indicates that short-term photodynamic stress primarily activates envelope stress response pathways, which aligns with membrane damage being an early and prominent target of photodynamic treatment [8], [27], [28], [29]. Additionally, Fur-controlled regulation likely plays a role in early control of intracellular iron levels, helping to limit Fenton chemistry under ROS-generating conditions [30]. Notably, standard oxidative stress regulons like OxyR and SoxRS were not significantly enriched, probably because induction of classic antioxidant genes occurs early (within 10 minutes) and wanes over time [19], while our RNA samples were taken after about 30 minutes of treatment, possibly after the peak of OxyR/SoxRS-driven transcription.

In contrast, long-term exposure preferentially engaged global transcriptional regulators, including ArcA, FNR, IHF, and NarL, indicating a transition from localized stress mitigation toward systemic metabolic and respiratory reprogramming. These regulators coordinate redox balance, anaerobic respiration, nitrogen utilization, and large-scale transcriptional organization under sustained stress conditions [31], [32], [33].

Overall, our data demonstrate that diverse photodynamic treatments elicit a shared acute stress architecture in *E. coli*, whereas prolonged exposure drives largely treatment-specific adaptive trajectories. The clear divergence between short- and long-term transcriptional responses highlights the importance of time-resolved analyses for separating immediate molecular damage from longer-term adaptive remodeling under photooxidative stress. Together, these findings reveal the remarkable transcriptional plasticity of bacteria in response to photodynamic challenge and provide a foundation for the rational refinement of light-based antimicrobial therapies.

## Acknowledgments

The authors have no acknowledgments to declare.

## Authors’ contribution

NB provided data and performed data analysis and interpretation. MWS supported data analysis and interpretation. MWS and MG contributed study set-up, evaluated and interpreted the data. NB and MG wrote the manuscript. All authors read and approved the final manuscript.

## Competing interests

The authors have declared no competing interests.

## Availability of data

The data generated or analyzed during this study are included in this published article and in the supplemental material. RNA-seq data are deposited in the NCBI Gene Expression Omnibus (GEO) database under accession number GSE317397.

## Notes

**Funding** This work was supported by the National Science Centre in Poland (Grant No. OPUS 2021/43/B/NZ6/01652 to MG).

## Bibliography

[1] WHO Bacterial Priority Pathogens List, 2024. 2024.

[2] R. Youf et al., “Antimicrobial photodynamic therapy: Latest developments with a focus on combinatory strategies,” Pharmaceutics, vol. 13, no. 12, pp. 1–56, 2021, doi: 10.3390/pharmaceutics13121995.

[3] A. Rapacka-Zdonczyk, A. Wozniak, M. Pieranski, A. Woziwodzka, K. P. Bielawski, and M. Grinholc, “Development of Staphylococcus aureus tolerance to antimicrobial photodynamic inactivation and antimicrobial blue light upon sub-lethal treatment,” Sci. Rep., vol. 9, no. 1, pp. 1–18, 2019, doi: 10.1038/s41598-019-45962-x.

[4] N. Kashef and M. R. Hamblin, “Can microbial cells develop resistance to oxidative stress in antimicrobial photodynamic inactivation?,” Drug Resistance Updates, vol. 31, pp. 31–42, Mar. 2017, doi: 10.1016/J.DRUP.2017.07.003.

[5] M. Lan, S. Zhao, W. Liu, C. S. Lee, W. Zhang, and P. Wang, “Photosensitizers for Photodynamic Therapy,” Adv. Healthc. Mater., vol. 8, no. 13, pp. 1–37, 2019, doi: 10.1002/adhm.201900132.

[6] Y. Wang et al., “Antimicrobial blue light inactivation of pathogenic microbes: State of the art,” Drug Resistance Updates, vol. 33–35, pp. 1–22, Nov. 2017, doi: 10.1016/j.drup.2017.10.002.

[7] F. Cieplik et al., “Antimicrobial photodynamic therapy–what we know and what we don’t,” Crit. Rev. Microbiol., vol. 44, no. 5, pp. 571–589, 2018, doi: 10.1080/1040841X.2018.1467876.

[8] E. Alves, M. A. F. Faustino, M. G. P. M. S. Neves, A. Cunha, J. Tome, and A. Almeida, “An insight on bacterial cellular targets of photodynamic inactivation,” Future Med. Chem., vol. 6, no. 2, pp. 141–164, 2014, doi: 10.4155/fmc.13.211.

[9] C. dos Anjos et al., “New Insights into the Bacterial Targets of Antimicrobial Blue Light,” Microbiol. Spectr., vol. 11, no. 2, Apr. 2023, doi: 10.1128/spectrum.02833-22.

[10] M. R. Hamblin and H. Abrahamse, “Oxygen-independent antimicrobial photoinactivation: Type III photochemical mechanism?,” Antibiotics, vol. 9, no. 2, pp. 1–17, 2020, doi: 10.3390/antibiotics9020053.

[11] D. Muehler et al., “Stress response in Escherichia coli following sublethal phenalene-1-one mediated antimicrobial photodynamic therapy: an RNA-Seq study,” Photochemical and Photobiological Sciences, vol. 23, no. 8, pp. 1573–1586, Aug. 2024, doi: 10.1007/S43630-024-00617-3/FIGURES/6.

[12] T. L. Adair and B. E. Drum, “RNA-Seq reveals changes in the Staphylococcus aureus transcriptome following blue light illumination,” Genom. Data, vol. 9, pp. 4–6, Sep. 2016, doi: 10.1016/j.gdata.2016.05.011.

[13] S. B. Snell, A. L. Gill, C. G. Haidaris, T. H. Foster, T. M. Baran, and S. R. Gill, “Staphylococcus aureus Tolerance and Genomic Response to Photodynamic Inactivation,” mSphere, vol. 6, no. 1, Feb. 2021, doi: 10.1128/MSPHERE.00762-20/SUPPL_FILE/MSPHERE.00762-20_ST003.XLSX.

[14] P. Walker et al., “Exploiting Violet-Blue Light to Kill Campylobacter jejuni : Analysis of Global Responses, Modeling of Transcription Factor Activities, and Identification of Protein Targets,” mSystems, vol. 7, no. 4, Aug. 2022, doi: 10.1128/msystems.00454-22.

[15] F. Ronzani et al., “Comparison of the photophysical properties of three phenothiazine derivatives: Transient detection and singlet oxygen production,” Photochemical and Photobiological Sciences, vol. 12, no. 12, pp. 2160–2169, 2013, doi: 10.1039/c3pp50246e.

[16] F. Harris and L. Pierpoint, “Photodynamic therapy based on 5-aminolevulinic acid and its use as an antimicrobial Agent,” Med. Res. Rev., vol. 32, no. 6, pp. 1292–1327, Nov. 2012, doi: 10.1002/med.20251.

[17] T. Baba et al., “Construction of Escherichia coli K-12 in-frame, single-gene knockout mutants: The Keio collection,” Mol. Syst. Biol., vol. 2, 2006, doi: 10.1038/msb4100050.

[18] V. Zorraquino, M. Kim, N. Rai, and I. Tagkopoulos, “The genetic and transcriptional basis of short and long term adaptation across multiple stresses in Escherichia coli,” Mol. Biol. Evol., vol. 34, no. 3, pp. 707–717, 2017, doi: 10.1093/molbev/msw269.

[19] M. Roth et al., “Transcriptomic Analysis of E. coli after Exposure to a Sublethal Concentration of Hydrogen Peroxide Revealed a Coordinated Up-Regulation of the Cysteine Biosynthesis Pathway,” Antioxidants, vol. 11, no. 4, 2022, doi: 10.3390/antiox11040655.

[20] S. A. Khan, M. J. Kim, and H. G. Yuk, “Genome-wide transcriptional response of Escherichia coli O157:H7 to light-emitting diodes with various wavelengths,” Scientific Reports 2023 13:1, vol. 13, no. 1, pp. 1–12, Feb. 2023, doi: 10.1038/s41598-023-28458-7.

[21] G. L. Müller et al., “Light Modulates Metabolic Pathways and Other Novel Physiological Traits in the Human Pathogen Acinetobacter baumannii.,” J. Bacteriol., vol. 199, no. 10, pp. 1–17, May 2017, doi: 10.1128/JB.00011-17.

[22] S. Jozefczuk et al., “Metabolomic and transcriptomic stress response of Escherichia coli,” Mol. Syst. Biol., vol. 6, no. 364, pp. 1–16, 2010, doi: 10.1038/msb.2010.18.

[23] G. Nonaka, M. Blankschien, C. Herman, C. A. Gross, and V. A. Rhodius, “Reveals a Multifaceted Cellular Response To Heat Stress,” Genes Dev., vol. 20, pp. 1776–1789, 2006, doi: 10.1101/gad.1428206.but.

[24] M. Zheng, X. Wang, L. J. Templeton, D. R. Smulski, R. A. LaRossa, and G. Storz, “DNA microarray-mediated transcriptional profiling of the Escherichia coli response to hydrogen peroxide,” J. Bacteriol., vol. 183, no. 15, pp. 4562–4570, 2001, doi: 10.1128/JB.183.15.4562-4570.2001.

[25] J. Courcelle, A. Khodursky, B. Peter, P. O. Brown, and P. C. Hanawalt, “Comparative gene expression profiles following UV exposure in wild-type and SOS-deficient Escherichia coli,” Genetics, vol. 158, no. 1, p. 41, 2001, doi: 10.1093/GENETICS/158.1.41.

[26] C. Janion, “Inducible SOS response system of DNA repair and mutagenesis in Escherichia coli,” Int. J. Biol. Sci., vol. 4, no. 6, pp. 338–344, 2008, doi: 10.7150/ijbs.4.338.

[27] S. Bury-Moné et al., “Global analysis of extracytoplasmic stress signaling in Escherichia coli,” PLoS Genet., vol. 5, no. 9, 2009, doi: 10.1371/journal.pgen.1000651.

[28] T. L. Raivio, S. K. D. Leblanc, and N. L. Price, “The Escherichia coli Cpx envelope stress response regulates genes of diverse function that impact antibiotic resistance and membrane integrity,” J. Bacteriol., vol. 195, no. 12, pp. 2755–2767, 2013, doi: 10.1128/JB.00105-13.

[29] G. Jovanovic, L. J. Lloyd, M. P. H. Stumpf, A. J. Mayhew, and M. Buck, “Induction and function of the phage shock protein extracytoplasmic stress response in Escherichia coli,” Journal of Biological Chemistry, vol. 281, no. 30, pp. 21147–21161, 2006, doi: 10.1074/jbc.M602323200.

[30] S. W. Seo, D. Kim, H. Latif, E. J. O’Brien, R. Szubin, and B. O. Palsson, “Deciphering Fur transcriptional regulatory network highlights its complex role beyond iron metabolism in Escherichia coli,” Nature Communications 2014 5:1, vol. 5, no. 1, pp. 4910–, Sep. 2014, doi: 10.1038/ncomms5910.

[31] S. Iuchi and E. C. C. Lin, “Adaptation of Escherichia coli to redox environments by gene expression,” Mol. Microbiol., vol. 9, no. 1, pp. 9–15, Jul. 1993, doi: 10.1111/J.1365-2958.1993.TB01664.X;WGROUP:STRING:PUBLICATION.

[32] D. A. Tolla and M. A. Savageau, “Regulation of Aerobic-to-Anaerobic Transitions by the FNR Cycle in Escherichia coli,” J. Mol. Biol., vol. 397, no. 4, p. 893, 2010, doi: 10.1016/J.JMB.2010.02.015.

[33] C. E. Noriega, H. Y. Lin, L. L. Chen, S. B. Williams, and V. Stewart, “Asymmetric cross-regulation between the nitrate-responsive NarX-NarL and NarQ-NarP two-component regulatory systems from Escherichia coli K-12,” Mol. Microbiol., vol. 75, no. 2, pp. 394–412, Jan. 2010, doi: 10.1111/J.1365-2958.2009.06987.X;PAGE:STRING:ARTICLE/CHAPTER.

